# Clinical, Biological and Pathological Investigation of Bovine Lymphocyte Intestinal Retention Deficit (BLIRD): a new genetic disorder affecting life expectancy and immunity in Holstein dairy cattle

**DOI:** 10.1101/2025.05.18.654695

**Authors:** Lucie Dutheil, Blandine Gausseres, Florian Besnard, Laurence Guzylack-Piriou, Yanad Abdou Monsef, Nicolas Gaide, Lisa Arnalot, Fabien Corbiere, Marie Gaborit, Frédéric Launay, Agnès Poujade, Aurélien Capitan, Gilles Foucras

**Affiliations:** Université de Toulouse, ENVT, INRAE, IHAP, 31076 Toulouse, France; Institut de l’élevage, 75012 Paris, France; Université Paris-Saclay, INRAE, AgroParisTech, GABI, 78350 Jouy-en-Josas, France; Ecole d’Ingénieurs de Purpan, Toulouse, France; GenPhySE, Université de Toulouse, INRAE, ENVT, Castanet-Tolosan, France; INRAE UE326, Unité Expérimentale du Pin, 61310 Le-Pin-au-Haras, France

**Keywords:** ITGB7, BLIRD, immune disorder, leucocytosis, lymphocyte, eosinophil, intestine

## Abstract

Dozens of missed recessive loci affecting homozygous carriers’ life expectancy were recently reported. This article details the clinical, biological and pathological manifestations of a new bovine genetic disorder caused by an ITGB7 p.G375S point mutation in the French Holstein cattle breed. Our thorough study involved database analysis of genotyped cattle and a series of case-control investigations of forty homozygous mutants for the causative variant. These mutants had a significantly shorter lifespan (fewer than 64% surviving past three years vs. 87% in controls), along with reduced body weight, daily weight gain, and dairy performance. The mutation did not affect most biochemical parameters, but a marked lymphocytic leucocytosis, moderate eosinophilia and differences in faecal microbiota were observed. Although non-pathognomonic symptoms may be confused with those of common environmental diseases, the blood profile effectively identified suspected carriers who developed ill-thrift and poor growth as heifers. Our research demonstrates that the bovine ITGB7 p.G375S substitution leads to reduced longevity, poor condition and production, in most homozygous carriers. Furthermore, this spontaneous model may help to refine the functions of the β7 integrin in immune homeostasis and defence.

## Introduction

Genetic selection (GS) has played a crucial role in advancing dairy cattle breeding, particularly in enhancing milk production, by the global use of a limited number of common dairy breeds such as Holstein and selected sires over the past century ^1^. This strategy has contributed to significant improvements in the genetic potential of dairy cattle. Additionally, the application of genomic information in bull breeding has enabled the selection of desired traits more effectively, while the integration of single nucleotide polymorphism (SNP) data has enhanced breeding efficiency by providing more precise predictions of genetic merit. While these advancements have led to remarkable gains in productivity, they have also raised concerns about the erosion of genetic diversity within dairy cattle populations. This reduction in genetic variation could undermine disease resistance and potentially shorten the lifespan of dairy cattle. Despite these breeding advances, the average lifespan of dairy cattle remains relatively short, typically between 4.5 and 5.5 years, with a productive life span of only 2.5 to 3.5 lactations, which is well below their expected life expectancy ^2,3^. Approximately 20% of dairy heifers do not survive to their first calving (USDA, 2007), and early culling, driven by factors like health problems and reproductive failure, leads to significant economic losses ^4^. The narrowing of genetic diversity within the Holstein breed has further compounded these concerns. As of 2017, the average genomic inbreeding level among Holstein bulls in the United States was 12.7%, with cows showing an inbreeding level of 7.9% ^5^. This inbreeding has negative effects not only on the longevity of dairy cattle but also on their production, fertility, and health traits ^6^. Moreover, the prevalence of genetic disorders is a growing concern, with early studies such as those by Robertson (1960) and Charlesworth and Charlesworth (1999), laying the groundwork for understanding the genetic basis of many inherited conditions. More recent studies, including those by Daetwyler et al., (2007) highlight the impact of modern breeding practices on the spread of genetic disorders. In addition to outstanding progresses in animal breeding and dairy production by identifying preferred traits (Brito et al., 2021) and predicting genetic merit ^11^, modern advancements in genomic technologies like GWAS and whole genome sequencing have significantly improved the detection of recessive mutations responsible for conditions like retinal degeneration ^10^ or Holstein haplotypes HH1, and HH2 ^12–16^. These mutations, identified by positional cloning and haplotype mapping, highlight the risks associated with intensive limited gene pool use ^17,18^. Recent research by Besnard et al. (2024) which uses haplotype enrichment and depletion, led to the discovery of 13 new recessive deleterious loci in Holstein cattle, many of which were linked to juvenile mortality, demonstrating the effectiveness of these initiatives in addressing genetic disorders. One particularly notable discovery is called “Bovine Lymphocyte Intestinal Retention Defect” (BLIRD), which is caused by a mutation in the gene encoding β7 integrin (ITGB7 p. G375S). This mutation has an allele frequency of 4.8% in the Holstein population with approximately 2.3 homozygotes per 1,000 individuals (Besnard, 2024). This deleterious substitution affects a residue that is completely conserved across 128 vertebrate orthologs ^19^ encoding for ITGB7, a member of a family of adhesion molecules. The protein forms a heterodimer with ITGA4 to create the α4β7 integrin complex (also known as LPAM-1), a key cell adhesion molecule expressed predominantly on activated or memory CD4+ T lymphocytes ^20^. This heterodimer plays a pivotal role in the trafficking of T cells from the bloodstream to the gastrointestinal tract ^21^, which is essential for maintaining intestinal immunity ^22^. α4β7 integrin is also constitutively expressed on naïve T and B cells at a relatively low level. Its expression is tightly controlled during lymphocyte activation and differentiation into effector and memory T cells ^23^. It is also expressed in Natural Killer (NK) cells, stimulated monocytes, macrophages and eosinophils ^21^. α4β7 is shared by several subtypes of leukocytes including lymphocytes ^24,25^, non-classical monocytes ^26^ and eosinophils ^27^. The α4β7 integrin complex is believed to be gut-specific due to its binding to Mucosal Addressin Cell Adhesion Molecule 1 (MAdCAM-1), which is exclusively expressed on intestinal endothelial cells and facilitates leukocyte extravasation into intestinal high endothelial venules ^28^. As suggested by preliminary pathophysiological analyses on a small cohort of cases and control Holstein cattle (Besnard et al. 2024) and data from the literature (Wagner, 1996), the replacement of a glycine with a neutral serine may affect integrin β7’s functions thereby disrupting the homing of immune cells to the gastrointestinal tract, and the maintenance of mucosal integrity. From a pathological perspective, the dysregulation of ITGB7 may contribute to the development of inflammatory diseases, characterized by aberrant immune cell trafficking to the gut, which is partly mediated by the α4β7 integrin ^30^. In view of these observations, the objective of our current study is to investigate the phenotypic, lesional, and immunological consequences of the p. G375S ITGB7 mutation in Holstein cattle. By conducting comprehensive analyses of large genotype and phenotype databases, on-field clinical examinations, necropsies, and laboratory tests, we present novel data elucidating the immune consequences and underlying pathophysiology of BLIRD.

## Materials and methods

### Outline of the participant’s attributes and equipment description

Blood samples were collected by licensed veterinarians, with owner consent for diagnostic testing. No animal was purposely bred for the study, and invasive sampling was performed post-mortem. All data were obtained with permission from French breeders and breeding organizations.

### Animals and records available from the French National Bovine Database

This study takes advantage of the large amount of genetic and phenotypic information recorded for management and selection purposes in the French Holstein population. We extracted from the French National Bovine Database the genotypes for the ITGB7 p. G375S substitution for a total of 740,276 females genotyped at a young age (typically within their first three months of life) with successive versions of the Illumina EuroGMD custom SNP array from 2019 to date as part of routine genomic evaluation (for details on the probe used, see Besnard et al. 2024). We also extracted a set of information on various life events (dates of birth, death, artificial insemination, calving, lactation start and end, and cause of death), as well as yield deviations corrected for environmental effects, computed within the framework of genomic evaluations for the females that started a productive career. This information allowed us to create the appropriate cohorts for each analysis performed in this study.

### Population-level survival analysis

The effect of the ITGB7 p. G375S substitution on juvenile mortality and premature culling was studied over a period of three years, corresponding to two years of rearing and one year of productive life in conventional French Holstein dairy farms. A total of 203,180 wild-type, 21,746 heterozygous, and 437 homozygous mutant individuals born at least three years before the analysis date were selected, and over the period considered, we calculated the daily counts and proportions of animals alive, dead by euthanasia or natural causes, or slaughtered for each genotype. Finally, at the end of the period, the proportion of live animals was compared pairwise between genotypes using Fisher’s exact test on the number of live animals versus the sum of dead and slaughtered animals.

### Population-level analysis of performance records

The effects of homozygosity and heterozygosity for the ITGB7 p. G375S substitution were estimated on 14 production, morphological and fertility traits routinely recorded for selection purposes. Yield deviation data, i.e., records adjusted for the non-genetic effects included in the models used for the official genomic evaluations carried out on behalf of the French breeding organizations were obtained from GenEval (for details on the models used, see https://www.geneval.fr/english) for cohorts of 38,751 to 247,206 wild-type, 3723 to 25,698 heterozygous, and 36 to 262 homozygous mutant females, depending on the trait (see **Supplementary Table 1** for details on traits and cohort sizes).

A mixed animal model including the fixed effects of the substitution (0 versus one or two copies) and year of recording, along with the individual random polygenic effect, was used to estimate the effects without bias Calculations were performed using blupf90+ from the *BLUPF90* suite of programs ^31^. To assess statistically significant differences between the group means of each trait, a Student’s t-test was applied using the stats R package. P-values were adjusted using the Benjamini-Hochberg method to control the false discovery rate ^32^. Finally, effects were standardized to genetic standard deviations (GSD) based on genetic parameters estimated from national genomic evaluations, to facilitate trait comparison.

In addition to yield deviations, we analyzed raw performances for three traits that are not considered in genomic evaluations but that we considered relevant for this study: age at first AI, which can be used as a proxy for growth in the absence of specific recording ^33^ (e.g., Jourdain et al., 2023), age at first calving, which is highly correlated with the previous one, and duration of the first lactation. The means between each genotype group were compared using a Student’s t-test.

### Retrospective analysis of heifer growth in an experimental farm

Monthly weight data recorded during the first two years of life were obtained from the INRAE experimental unit at Le-Pin-au-Haras (Orne, France) for 719 wild-type, 131 heterozygous and 4 homozygous mutants for the ITGB7 p. G375S substitution. The growth curves of each animal were determined using regression. Then, the analyses performed included the determination of the curves corresponding to the 10th and 90th percentile and the mean curves for the wild-type and heterozygous genotypes. Due to the small number of homozygous mutants and the fact that they all died before reaching the end of the rearing period, individual growth curves were displayed for this group.

### Field survey

A field survey was conducted on female cattle homozygous for the ITGB7 G375S substitution to evaluate further the point mutation impact drawn from population records. From a list of 314 live homozygote females distributed throughout France, 40 cattle were selected in 38 commercial herds distributed across Normandy and the Southwest region of France. The study’s methodology involved the choice of a matched control in the same herd with the lowest age difference, in most cases less than one month. Following prevailing farm practices, the animals commonly received preventive treatments, including respiratory vaccines, deworming drugs, and anticoccidials. However, these treatments were not the subject of evaluation in the present study. In total, 86 animals were included in the survey and subjected to a close clinical examination (40 homozygous mutants, 10 heterozygotes and 36 wild-type variant carriers). Each animal was clinically examined and photographed alongside its control. Live weight was indirectly assessed through the thoracic circumference using a barometric ribbon, and body weight was deduced using the Crevat formula (P = K T^3^, where T is the chest circumference in metres, K is a coefficient that has been calculated on average at 80). Three of the homozygotes included in the field survey were brought to the ruminant clinic of the National Veterinary School of Toulouse (ENVT) for detailed clinical examination and tissue sampling after necropsy.

### Laboratory analysis

Blood samples were collected from the 86 animals of the field survey for subsequent serum analysis and complete blood count. Venous blood sampling was conducted in Vacutainer® EDTA, heparin or dry collection tubes. The blood samples were stored at 4°C, and processed within 12h by centrifugation for serum and plasma separation. Samples collected in EDTA tubes were used for haematological purposes, as heparin and dry tubes were used for biochemistry and serology respectively.

### Biochemistry

For biochemistry, the following parameters were assessed on heparin anticoagulated plasma: total proteins, albumin, urea, and aspartate aminotransferase (ASAT) in 54 animals (half mutants and half controls). Globulins were calculated as the difference between total protein and albumin. Analyses were conducted using a Vitros 250 analyser and appropriate multi-layer reagents (Ortho Clinical Diagnostics, Issy les Moulineaux, France), following the manufacturer’s instructions. Quality control of the Vitros 250 was performed using the Performance Verifier I & II solutions (Ortho Clinical Diagnostics, Issy les Moulineaux, France).

### Complete blood count

Prior to analysis, the EDTA blood tubes were held at room temperature for 20 minutes, during which time they were gently agitated to ensure homogenisation. Measurements were conducted within a 12-hour window after sampling, carried out by the CREFRE – Inserm – UPS – ENVT Comparative Medical Biology and Histology platform using Sysmex XN-V and Sysmex XT-2000iV software. Blood cell enumeration was also done using air-dried blood smears and staining with a May-Grünwald/Giemsa automatic stainer. A comprehensive range of variables was analysed, including impedance and optical red blood cell (RBC) counts, haematocrit, haemoglobin concentration, mean corpuscular volume, red cell distribution, platelet counts, mean platelet volume, platelet crit, platelet distribution width, and platelet large cell ratio. The variables measured by the analyser include the white blood cell (WBC) count, as well as neutrophil, lymphocyte, monocyte, eosinophil and basophil counts. These were determined from 100 leukocytes counted per oil immersion field ^34^.

### Flow cytometry analysis

Flow cytometry was conducted on mononuclear cell fractions from blood, mesenteric lymph node (mLN) and lamina propria (LP) of homozygous mutants (n=3) and non-carriers (n=5). PBMC were prepared from EDTA-anticoagulated whole blood diluted 1:1 with phosphate-buffered saline (Dutsher, Cat#L0615), layered on half the volume of Ficoll Paque Plus (Cytiva, Cat#17-1440-03) and centrifugated at 1200 xg for 20 min at room temperature with the brake off. Red blood cells were lysed with ACK lysis buffer (154 mM ammonium chloride 10 mM potassium bicarbonate, 97 µM EDTA, pH 7,4). mLN were smashed using a syringe piston and a 70 µm cell strainer to produce a cell suspension.

The small intestinal (jejunum) wall was washed in cold PBS, cut into pieces, and incubated four times with 3 mM EDTA for 40 minutes at 37 °C. Tissue pieces were then rinsed and digested in DMEM with 20% FCS and 100 U/ml of collagenase for 40 minutes further at 37 °C. The suspension was filtered and undigested pieces were smashed on a 70 µm cell strainer using a syringe piston. Finally, mononuclear cells were isolated using a 40–80% Percoll gradient.

Cells were resuspended in HBSS with 0.5% BSA and 10 mM Hepes. Cell viability was assessed using Viobility 405/520 Fixable Dye (Miltenyi Biotec, Cat#130-130-404). The antibodies used were: CD45 FITC (BioRad, Cat#MCA2220), CD4 Pacific Blue (BioRad, Cat#MCA1653), CD45RO Alexa Fluor 647 (BioRad, Cat#MCA2434), α4 integrin PE/Cyanine7 (BioLegend, Cat#304314) and β7 integrin PE (BioLegend, Cat#121006).

Antibody incubation was performed for 20 minutes at 4°C in the dark. Data were collected on a MACSQuant® Analyzer (Miltenyi Biotec) and subsequently analysed using FlowJo software (Becton Dickinson).

### Faecal microbiota composition

To assess the composition of the gut microbiota, faeces were collected directly from the rectum and placed in a leak-proof plastic container. Bacterial microbiota from 48 samples (24 homozygous mutants and 24 wild-type controls) was investigated as described by Arnalot, (submitted). Briefly, DNA was extracted from faecal samples using the Quick-DNA Faecal/Soil Microbe Miniprep Kit (Zymo Research, Irvine, CA, United States of America) according to the manufacturer’s instructions. The 16S rRNA V3-V4 region was investigated and the sequencing was performed using MiSeq Illumina Sequencing at the Genomic and Transcriptomic Platform (INRAE, Toulouse, France).

The sequenced reads were analyzed using FROGS v4.1.0 software (Escudié et al., 2018), with parameters described in (Arnalot et al submitted). Briefly, 1,709,057 raw sequences were pre-processed, resulting in 1,492,612 sequences retained (87.3%), with the number of reads per sample ranging from 23,552 to 39,187. A total of 2,173 clusters and 1,196,762 sequences were retained, with a range of 18,456 to 32,470 sequences per sample. The subsequent data were computed in a *phyloseq* object for further R analysis (McMurdie and Holmes, 2013).

### Serological testing and pathogen detection

Both serological assays and pathogen detection through polymerase chain reactions (PCR) were employed on 21 couples in the field study cohort. Diagnostic tests were performed using commercially available kits from ID vet Genetics and ID Screen, following the manufacturer’s protocols. A duplex PCR assay (ID Gene Paratuberculosis Duplex, ID vet Genetics) was used to detect *Mycobacterium avium* subsp. *paratuberculosis* (Johne’s disease) on DNA extracted from faecal samples. For the concurrent detection of Bovine Viral Diarrhea Virus (BVDV), a triplex PCR assay (ID Gene BVDV/BDV Triplex 2.0, ID vet Genetics) was used, with blood-extracted RNA serving as the test matrix. In addition, serological screening for Bovine herpesvirus I and BVDV antibodies was conducted using enzyme-linked immunosorbent assays (ELISA), with ID Screen IBR gB Competition (Innovative Diagnostics, Grabels, France) to detect antibodies against the gB Bovine Herpesvirus type 1 (BoHV-1) glycoprotein and ID Screen BVD p80 Antibody Competition (Innovative Diagnostics, Grabels, France) for BVDV against the NS3 protein, with serum as the matrix. For Paratuberculosis, an indirect ELISA (ID Screen Paratuberculosis Indirect) was also used to detect antibodies against *M. avium* subsp. *paratuberculosis* in bovine serum. qPCR analysis was done on DNA faecal extracts to quantify the presence of Ostertagia/Cooperia eggs in the faeces of select animals ^35^. All diagnostic assays were conducted in strict accordance with the manufacturer’s instructions.

### Post-mortem assessment & histopathology

A complete necropsy was performed after euthanasia on three homozygous mutant heifers, 28 months-old (n=2) and 17 months-old (n=1). For histopathology section of heart, liver, spleen, thymus, lung, rumen, abomasum, duodenum, jejunum, ileum, caecum, proximal and spiral colon, rectum, laryngeal and lingual tonsils, as well as peripheral and loco-regional visceral lymph nodes (left ruminal, ventral abomasal, jejunal, caecal, cranial mesenteric and caudal mesenteric, left mandibular, superficial cervical, subiliac, deep popliteal, iliofemoral, left tracheobronchial) were collected. All samples were fixed in 10% neutral buffered formalin for histopathological examination and immunohistochemistry. In parallel, intestinal sections (ileum) and the draining lymph node from five healthy homozygous wild type Holstein female cattle were collected at the slaughterhouse to provide a histological comparison of tissues within normal limits.

Fixed tissue samples were paraffin-embedded and cut at 3 µm. Sections were stained with haematoxylin and eosin (H&E) for histopathological analysis. Immunohistochemistry (IHC) was performed to determine the distribution and abundance of lymphocytic B and T populations in the jejunum, jejunal lymph node and prescapular lymph node tissues, using CD3 (T-lymphoid cell marker, mouse monoclonal, Clone F7.2.38, Agilent, 1/50, M 7254) and CD20 (B-lymphoid cell marker, rabbit polyclonal, Thermo scientific, 1/600, PA5-16701). Briefly, the IHC protocol included an antigen retrieval step with buffer solution (EnVision Flex Target Retrieval Solution, Low and High pH for CD3 and CD20 respectively, Agilent) applied for 30 min at 96°C, a peroxidase blocking step of 5 min at room temperature (S2023; Agilent) followed by saturation of nonspecific binding sites with normal goat serum (X0907; Agilent) applied for 20 min at room temperature, and primary incubation for 50 min at room temperature. Signal amplification and revelation were assessed using the EnVision FLEX system and 3,3-diaminobenzidine (DAB) revelation according to the manufacturer’s recommendations.

### Statistical analysis of the data from the field survey cohort

Univariate comparisons of all measured parameter distributions between groups were performed using the Mann-Whitney-Wilcoxon test for ranks. The heterozygotes and wild-type homozygotes were merged into a single control group (ctrl) due to lack of significant difference between them, and compared with the mutant homozygote group (mut). All the results are provided in **Supplementary Table 5**. Data visualization and statistical analysis were performed using R software ^36^.

Generalised linear mixed models (GLMMs) were used for multivariate analysis, with response variables comprising morphological or biological parameters, including chest circumference, haemoglobin levels, lymphocyte count, monocyte count, eosinophil count, basophil count, urea levels, globulin levels and aspartate transaminase levels. Explanatory variables included the genotype group, age and farm. The *glmmTMB* package ^37^ was utilised to fit a series of models with varying degrees of complexity. The study considered various models, including simple models with fixed effects of Genotype and Age. The investigation also included the implementation of more intricate models that utilised natural B-splines to ensure the smoothness of the relationship between the quantitative variables and the outcome. The examination involved the interaction between Genotype and Age, models with or without random Farm effects, and where appropriate, models incorporating random farm effects and an interaction between genotype and age, as well as potential interactions between all three factors. The set of candidate models was fitted for each response variable, exploring various combinations of fixed and random effects. The most suitable model was selected based on the Akaike Information Criterion (AIC). The *emmeans* ^38^ package was used to calculate estimate the marginal effects of genotype at different ages, with Age in models that included it as a fixed factor in the models. Marginal estimates were calculated at different ages and compared between genotype groups.

Multivariate relationships among continuous variables were examined using multiple factor analysis (MFA) and principal component analysis (PCA). Hierarchical clustering on principal components (HCPC) was then performed to assess genotype robustness. MFA, PCA, and HCPC were performed using the *FactoMineR* package ^39^.

Partial least squares discriminant analysis (PLS-DA) and sparse partial least squares discriminant analysis (sPLS-DA) were used to improve the ability of the model to discriminate linear combinations of variables and to identify the most influential variables in genotype classification via the *MixOmics* package ^40^. This supervised method reduces the dimensionality of the data and has been shown to provide superior results in terms of improved genotype separation. This technique helps to discriminate between two sets of labelled points by identifying the most influential variables in genotype classification.

Regarding faecal bacterial community, *adonis2* from the *vegan* package ^41^ was used to test for β-diversity significance between groups. The significant differential taxa were tested with Analysis of Compositions of Microbiomes with Bias Correction 2 (*ancom-bc2,* package *ANCOMBC* ^42,43^).

Statistical significance was defined as a p-value less than 0.05, and the results meeting this criterion were considered significant.

## Results

### Retrospective analyses indicate that the ITGB7 p.G375S substitution is associated with reduced longevity and poor zootechnical performance

To gain insight into the consequences of the ITGB7 p. G375S substitution on bovine health and longevity, we first studied the survival over three years of a large cohort of females (n= 437 homozygous mutants (hmz), 21,746 heterozygous (htz) and 203,180 wild type (wt) animals) genotyped for the corresponding DNA variant as part of the routine genomic evaluation. Survival, mortality and slaughter curves are shown in **Figure 1A**.

**Figure 1:**
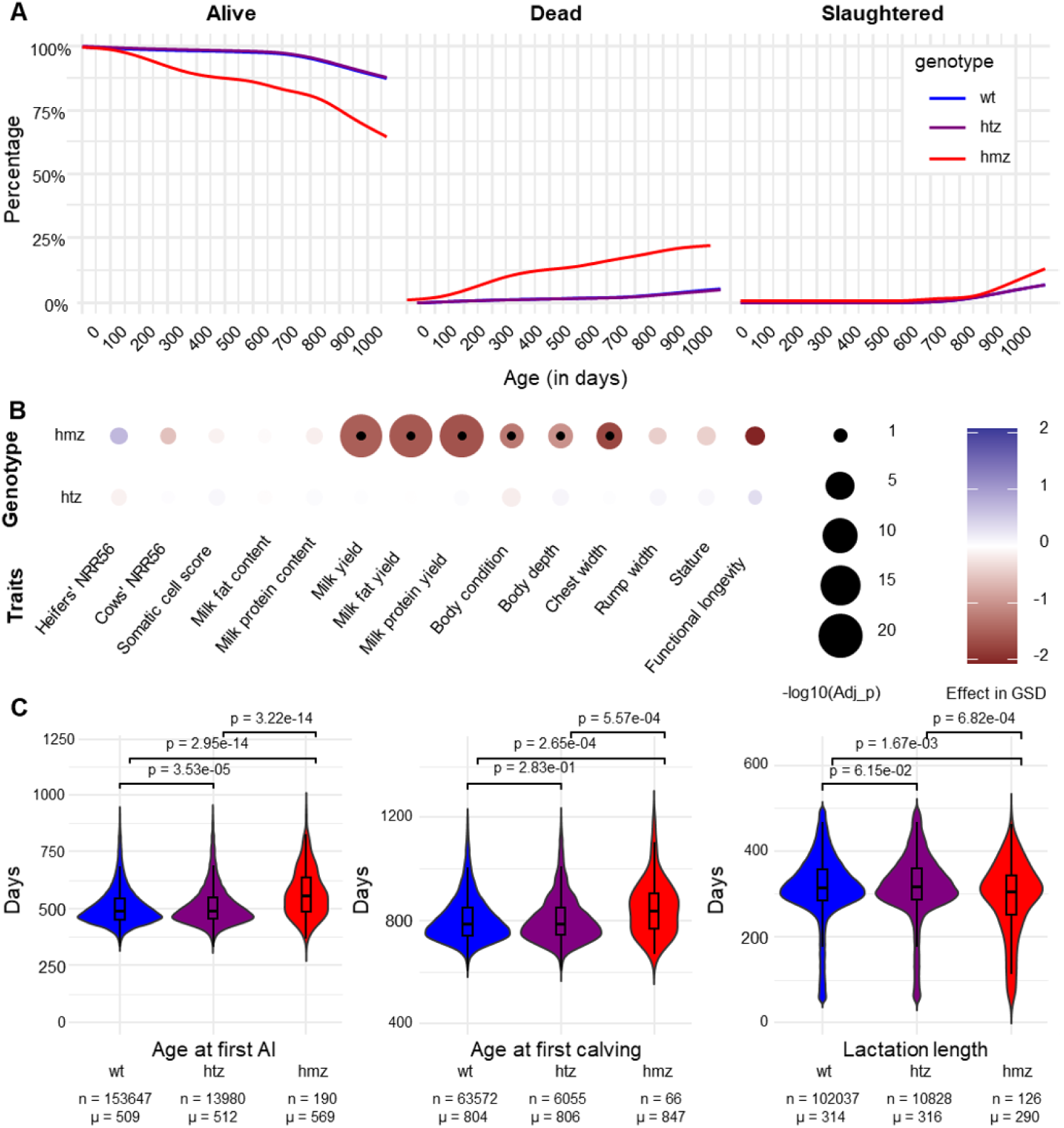
Population scale analyses of survival and performance records for Holstein females genotyped for the ITGB7 p. G375S substitution. (**A**) Proportion of animals alive, dead (by euthanasia or natural causes), or slaughtered for 203,180 wild-type, 21,746 heterozygous, and 437 homozygous mutant individuals over three years, corresponding to two years of rearing and one year of productive life in conventional French Holstein dairy farms. (**B**) Effects of homozygosity and heterozygosity for the ITGB7 p. G375S substitution on 14 traits expressed in genetic standard deviation (GSD; see Supp Tab <u>1</u> for details on cohort sizes). –log10(Adj-p): – log10 of Benjamini–Hochberg adjusted Student’s t-test p values. (**C**) Analysis of age at first AI, age at first calving and duration of first lactation. Means and cohort sizes are shown below the violin plots for each genotype group. The p-values shown for each comparison were obtained using a Tuckey test.

The red curve corresponds to mutants (mut); meanwhile, the blue and violet curves correspond to heterozygotes (htz) and wild-type homozygotes (wt). The curves for these two categories overlap. On the other hand, homozygous mutants showed a significant reduction in the proportion of live animals at the end of the period with only 63.6% versus 87.5% for the heterozygotes, and 87.4% for the wild-type individuals (Fisher’s exact test p-values = 3.13 x 10^−36^ and 7.90 x 10^−37^, for each comparison respectively; **Supplementary Table 2**). This difference was due to a continuous excess due to either natural death and/or early slaughter between 2.5 and 3 years.

To investigate the causes of this increase in premature culling, we analysed yield deviations (i.e., phenotypes adjusted for environmental effects) for 14 traits considered in the French genetic evaluations ^19^, using a mixed model including the fixed effect of the ITGB7 substitution (0 versus 1 or 2 copies), the fixed effect of the year of recording, and the individual random polygenic effect (**Supplementary Table 1; Figure 1B**). The homozygous mutants appeared to have normal fertility, as measured in heifers (nulliparous) and cows (multiparous) using the 56-day nonreturn rate (NRR56, which considers the proportion of females that have not been re-inseminated 56 days after previous insemination and are considered pregnant). However, those showed significantly lower performance in several production and morphological traits. Homozygosity for the ITGB7 p.G375S substitution had no effect on milk composition (i.e. fat and protein content) but a strong negative effect on quantity, reducing milk yield by 1359 kg (equivalent to –1.79 genetic standard deviation or GSD; Benjamini Hochberg adjusted Student’s t-test p-value (FDR) = 3.30 × 10⁻^22^), milk fat yield by 58.02 kg (–1.86 GSD, FDR = 4.57 x 10^−24^) and milk protein yield by 43.87 kg (–1.93 GSD, FDR = 5.87 x 10^−24^). Highly negative significant effects in homozygotes were also observed (**Figure 1B**) for body condition (–1.50 GSD, FDR = 6.74 x 10^−4^) body depth (–1.21 GSD, FDR = 8.01 x 10^−4^) and chest width (–2.07 GSD, FDR = 4.74 x 10^−5^). Of note, suggestive but not significative effects after accounting for multiple testing were also observed in homozygous for stature (–0.51 GSD, raw p-value (raw-p) = 0.04) and functional longevity (–2.45 GSD; raw-p=0.02). Therefore, the homozygous mutants with available yield deviations are those less affected by the BLIRD. To further investigate the effects of the homozygosity for the ITGB7 p. G375S substitution on larger and less censored populations we studied raw phenotypes for three traits (**Figure 1C**). Age at the first insemination was significantly delayed by approximately 2 months in the homozygous mutants compared to the other genotypes (mean = 569 versus 512 and 509 days for the heterozygous and non-carriers, respectively; Tukey test p-value = 3.22 x 10^−14^ and 2.95 x 10^−14^ for the comparisons with each group respectively). Similar results were obtained for the age at first calving which is highly correlated to the latter trait in the absence of fertility problems. Finally, in concordance with the higher rates of premature culling observed in the survival analysis, the duration of the first lactation was also reduced by three to four weeks on average in the homozygous mutants as compared with the other genotypes (mean = 290 versus 316 and 314 days for the heterozygous and non-carriers, respectively; Tukey test p-value =6.82 x 10^−4^ and 1.67 x 10^−3^ for the comparisons with each group respectively).

To further investigate this effect on heifer growth, we analysed the monthly weight data of 719 wild-type, 131 heterozygous and 4 homozygous mutant heifers for the ITGB7 p. G375S substitution routinely recorded during their first two years at the INRAE experimental unit located at Le-Pin-au-Haras (Orne, France). The results showed (**Figure 2A)** that the growth curves of the heterozygous (purple curve) individual were not different from those of animals of the wild genotype (blue curve). However, the four homozygotes (in red) exhibited a growth curve that consistently fell below the lower tenth percentile of the farm’s growth curves during the period considered. Moreover, three cases died prematurely at 351, 468 and 653 days of age without prodromal signs.

**Figure 2:**
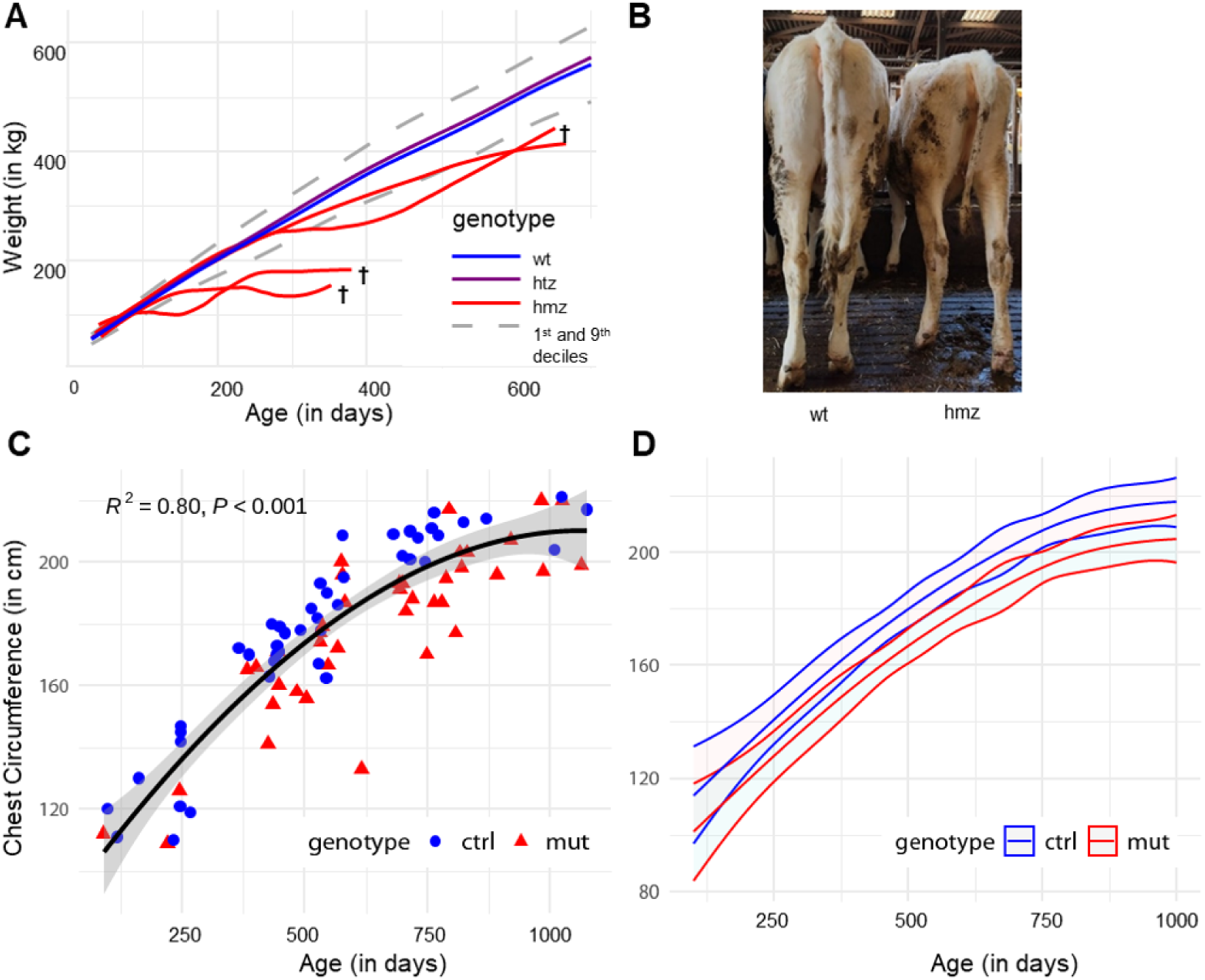
Growth data of ITGB7 p.G375S mutants (n=40) and controls (n=46). (**A**) Weight growth curves for heterozygous (htz), wild-type homozygous (wt), and mutant homozygous (mut) animals over 0-730 days in a single experimental herd. The graph presents the average weight curves for each genotype (mut, wt, and htz), along with the 10th and 90th percentiles derived from the total dataset. Individual weight curves are also displayed for the 4 mutant homozygous females (mut). (**B**) The ill-thriving phenotype of a homozygote mutant and its age-matched wild-type control, with less than 1-month difference. (**C**) Measured chest circumference (in cm) across age (days) for mutants (red triangle) and wild-type (blue round) animals. (**D**) Modelled impact of the ITGB7 p.G375S substitution on chest circumference over time according to the mutant or wt homozygous genotype.

### Growth retardation and ill-thrift in p.G375S ITGB7 homozygote carriers on-field

The 86 animals’ field survey revealed no notable differences between the 10 heterozygous and 36 wild-type heifers (Mann-Witney, p-value < 0.05), in any of the examined traits, confirming observations on Nation-wise records. As a result, the heterozygous cows were considered suitable as controls and were included in the control group (Ctrl). Later on, wild-type and heterozygous animals form the control group.

According to the results of the clinical examination on farms, many cases displayed reduced stature, sparse coats, and occasionally a larger abdomen. Despite exhibiting signs of ill thrift, no diarrhoea or loss of appetite were reported. **Figure 2B** depicts one mutant (mut) and its matching control (ctrl). The average chest circumferences and deduced weight of the mutants were not different from those of the controls (178±27 VS 178±31 cm, Mann-Whitney test, p-value=0.74). However, a graphical representation of chest circumference (in cm) according to the age (in days), **(Figure 2C**), reveals two discernible clusters. A pattern emerges in which animals with the mutation fall below the curve representing the third-degree polynomial regression of the Holstein breed standards ^44^. In contrast, the controls tend to be above or on the curve, irrespective of their age at the recording date, even though the mutant group is significantly older than the control group (642±225 vs 536 ± 234 days, Mann-Whitney test, p-value=0.03). Significant effects of genotype, age and farm on the response variable were evidenced by models and were accounted for to provide the most accurate estimate of the marginal effect of the genotype group (**Supplementary Table 3**).

The optimal model for predicting chest circumference, based on the lowest AIC (657.4), BIC (679.5), log-likelihood (–319.7), deviance (639.4), and 77 residual degrees of freedom, balances simplicity and fit. It uses a Gaussian family with an identity link and includes “genotype,” “age,” and a random effect for “farm.” Random effects analysis showed farm-level intercept variance of 80.63 (SD = 8.98), indicating variability in chest circumference across farms. Fixed effects revealed that the “mutant” genotype was associated with a significant reduction in chest circumference (–13.10 ± 1.75 cm, p = 6.94e-14), independent of age. Age, modelled by natural splines, showed a non-linear effect, with all spline terms significantly positive (p < 2e-16), and the largest effect at the fourth spline (147.355). In conclusion, this model highlights the significant impact of the G375S mutation on chest circumference, with genotype and age as key factors, and a random farm effect incorporated. Consequently, the model indicates ITGB7 p. G375S homozygotes is associated with reduced chest circumference. The estimated marginal means (EMMs) for each genotype is visualized in **Figure 2D** and various chosen age points are provided in **Supplementary Table 4**. Notably, for chest circumference, the mutated animals in the model would have a chest circumference at 480 days of 164 cm VS 177 cm for the controls. The calculated weight around breeding time (which corresponds to 356 kg VS 441 kg) representing only 50% of the desired mature weight of 700 kg, which may explain the delays in breeding the mutants.

The analysis of variables in models shows significant genotype effects on several variables, including chest circumference, haemoglobin, lymphocytes, monocytes and eosinophils. In particular, chest circumference showed a strong Genotype effect (chi-squared = 68.61, p < 1.20e-16) and a significant non-linear Age effect (chi-squared = 258.91, p < 6.72e-54). However, the genotype by age interaction for chest circumference was not significant (p = 0.07). Genotype was also a significant (p <0.05) factor for haemoglobin, lymphocytes, monocytes and eosinophils (**Supplementary Table 3**). In addition, age had a significant effect on urea (chi-squared = 26.56, p = 6.94e-05).

### ITGB7 p.G375S homozygote carriers have a severely perturbed haematological profile with a severe lymphocytosis

BVDV and BoHV-1 molecular and serological detections were negative for both cases and controls, except in a few vaccinated herds to BVDV, supporting the absence of active and ancient infection. Similarly, *Mycobacterium avium* subsp *paratuberculosis* faecal detection was all negative, and they were serologically negative for *M. paratuberculosis* antibodies, excluding a major role of these infections in the clinical findings.

To assess the role of ITGB7 p.G375S substitution on inflammatory parameters, systemic biomarkers in blood and blood biochemistry were evaluated between genotype groups **(Supplementary Table 5**). Urea levels were not different between the control and mutant groups (3.6±1.4 vs. 3.0±1.5 mmol/L respectively, Mann-Whitney, p=0,14). Similarly, aspartate aminotransferase (AST) values did not vary according to the group (118±127 vs. 104±40, Mann-Whitney, p=1). Plasma total proteins (73.5±8.7 vs. 75.3±6.5 g/L, Mann-Whitney, p=0,11), and albumin concentrations (33.6±5.7 vs. 31.3±3.4 g/L p=0,13) were not different on average, while globulinemia was significantly higher in the mutant group (39.9±4.0 vs. 44.0±4.5 g/L, Mann-Whitney test, p=0,001).

In contrast to biochemistry, complete blood count (CBC) exhibits variation according to the genetic group. Mean white blood cell count (WBC) was 150% higher (15.7±3.7 vs 10.2±2.3×10⁹, Mann-Whitney test, p-value<0.001) in the mutant group **(Figure 3A)**, and a significant difference was observed for lymphocyte count, (8.2±2.4 x 10⁹ vs 5.2 ± 1.6 x 10⁹, Mann-Whitney test, p-value<0.001) (**Figure 3B**). A markedly elevated population of eosinophils is present in the homozygote carriers in comparison to the control group (0.6±0.6 vs 1.4±0.7, Mann-Whitney, p<0.001). Similarly, monocyte count (2.2±1.1 vs 1.0±0.7, Mann-Whitney, p<0.001) and basophils (0.11±0.06 vs 0.08±0.08, Mann-Whitney, p<0.001) were also higher in the former contributing to the CBC difference. In contrast, the neutrophil counts (3.9±1.9 vs 3.3±1.4×10^9^ cell/L, Mann-Whitney test, p =0,11) and the red blood cell count (RBC, 7.2±1.0 vs 7.7±1.0×10^10^ cell/L, Mann-Whitney test, p =0,03), were within the reference range and not different between the groups. haemoglobin concentration (Hb, 10.4±1.4 vs 11.1±1.3 g/L, p =0,005), was significantly lower in the mutant group. A comprehensive overview of the biochemical and haematologic findings between controls and mutants is provided in **Supplementary Table 6**.

**Figure 3:**
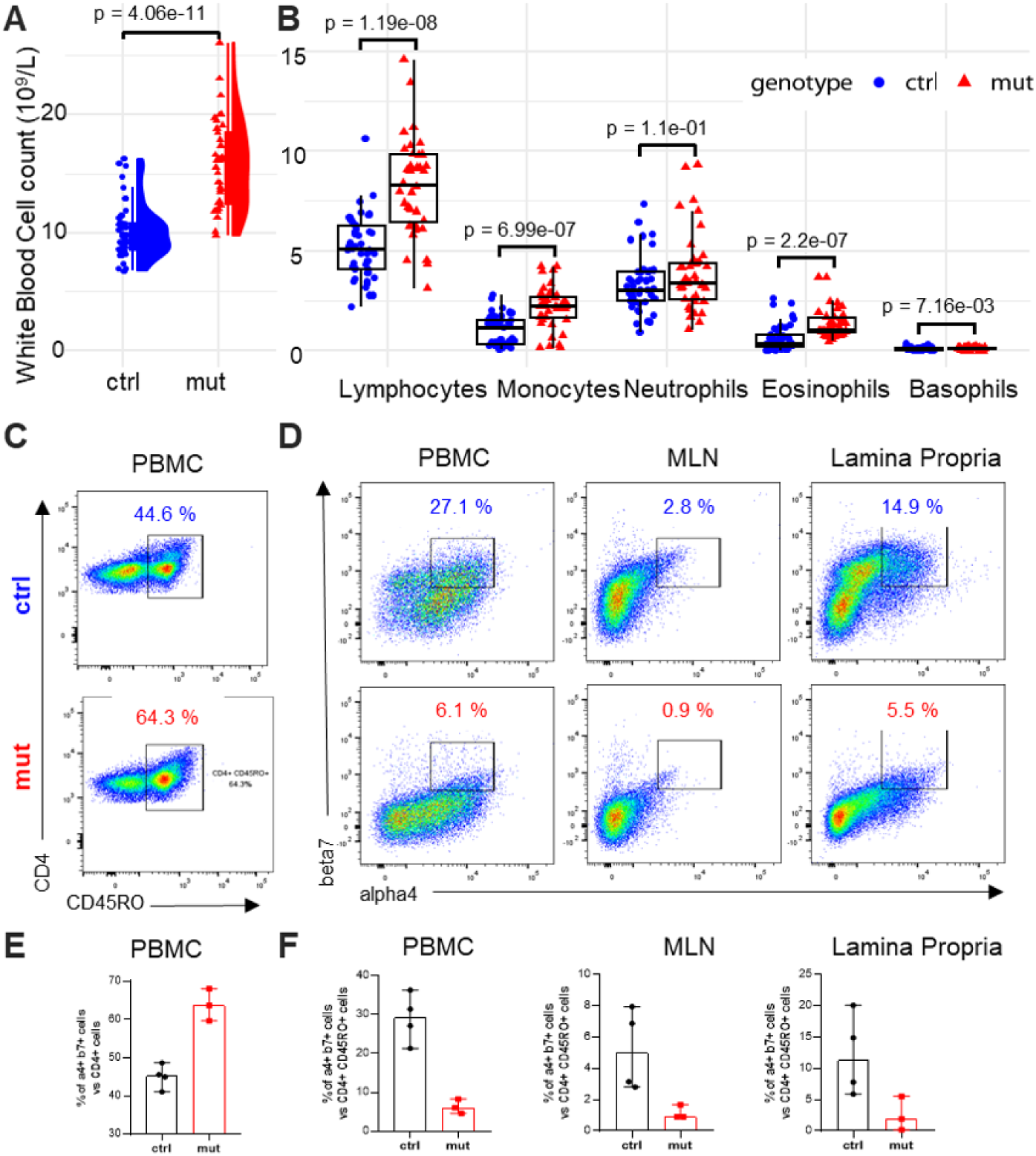
Blood count and tissue distribution of LPAM1^pos^ CD4 T cells in mutant and control animals. (**A**) White blood cell count (in 10^9^ cells/L) in ITGB7 p.G375S mutants and wild-type animals. (**B**) Blood Cell counts (in 10^9^ cells/L) in lymphocytes, monocytes, neutrophils, eosinophils, and basophils in mutant (n=36) and control (n=40) animals. (**C**) Representative plots of flow cytometry analysis for memory (CD45RO^pos^) CD4+ T lymphocytes in PBMC samples from three ITGB7 p.G375S homozygous mutants and five controls. (**D**) Representative plots for flow cytometry analysis of α4p^os^β7^pos^ double-positive cells among memory CD45RO^pos^ CD4^pos^ T lymphocytes in the blood (PBMC), mesenteric lymph nodes (MLN) and lamina propria (LP) samples of one homozygous mutant and one control. (**E**) Proportion of memory CD45RO^pos^ lymphocytes among CD4^pos^ T lymphocytes in PBMCs from ITGB7 p.G375S mutants (n=3) and wild-type controls (n=5). (**F**) Proportion of α4p^os^β7^pos^ double-positive cells among memory CD45RO^pos^ CD4^pos^ T lymphocytes in the blood (PBMC), mesenteric lymph nodes (MLN) and lamina propria (LP) samples of homozygous mutant (n=3) and homozygous control (n=5).

We subsequently examine the lymphocyte population to look for differences in their distribution across blood and digestive tissues, and to see how they can explain the difference in lymphocyte count between the two groups. LPAM-1 (ITGA4/ITGB7 dimer) has been shown to mediate the homing of memory (recognized as CD45RO^pos^ in cattle) CD4 T lymphocytes into the gut area (Wagner, 1996; William 1997). Naive T cells exhibit low levels of expression of CD45RO, which increase upon activation and differentiation into effector and memory T cells (Kilshaw & Murant, 1991). For that reason, we examined the ITGA4 and ITGB7 expression on CD45RO^pos^ CD4^pos^ T cells by flow cytometry in the blood on a small group of animals (three homozygous mutants and five controls). **Figures 3C** and **3E** shows that the mutant cows have a larger memory cells population among CD4 T lymphocyte than the control animals (64.3% VS 44.6%). Furthermore, the proportion of cells expressing ITGA4/ITGB7 among CD45RO^+^CD4^+^ T cells was reduced in homozygous animals compared to controls (**Figure 3D**); either in the blood (6.1% VS 27.1%), in the mesenteric lymph nodes (0. 9% VS 2.8%) and in the intestine (jejunum) of the mutants, which contains 2.7 times less LPAM-1^pos^ lymphocytes compared to the controls (5.5% VS 14.9%), as shown in **Figure 3F**.

These findings support the idea that BLIRD cattle have an impaired retention of CD4 T lymphocytes in the LP, and an observation behind the BLIRD designation of the genetic disorder, with fewer T cells ITGA4/B7 T cells in mesenteric lymph nodes (mLNs), and submucosal space (LP), despite overabundant lymphocytes in the blood.

### Histopathological findings

At necropsy, mutants weighed 300, 387 and 550 kg. All examined organs were within normal limits with no evident abnormalities that could explain the animal’s suboptimal health conditions or premature demise. Histologically, mutated animals presented moderate to marked cortical follicular lymphoid hyperplasia in peripheral (prescapular) lymph nodes (3/3) and intestinal payer’s patches (3/3) (**Figure 4A**, Hemalun&eosin staining). Paracortical areas were within normal limits or mildly reduced in size due to expansive areas of follicular hyperplasia in mutated animals. Other lymphonodal follicular changes were observed including intra-follicular haemorrhages, hyalinosis and dystrophic mineralisation. Intestinal mucosa as well as other tissues were within normal limits in comparison with control tissue from wild-type subjects. Immunohistochemistry confirmed the abundance of CD20^pos^ lymphoid cells within affected lymph nodes and Peyer’s patches, with prominent follicular organization (**Figures 4B & C**). CD20^pos^ cells frequently showed a moderate (2/3) to marked (1/3) increase in number and frequency within intestinal lamina propria in mutant subjects. In contrast, CD3^pos^ lymphoid cells were present, heterogeneously distributed throughout paracortical areas of the lymph nodes from mutant subjects. They were less frequent and mildly reduced in density in the jejunal lymph node (3/3) and prescapular lymph node (2/3). In intestinal mucosa, CD3^pos^ cells were observed expanding heterogeneously in the deep lamina propria of mutant subjects (3/3), in contrast with tissues from the controls where CD3^pos^ cells were mostly observed within superficial villous interstitium and epithelium. Overall histological data suggest intestinal, peripheral and loco-regional lympho-nodal B-cell lymphoid hyperplasia. No significant changes in T-cell density were observed but substantial topographical changes were noted.

**Figure 4:**
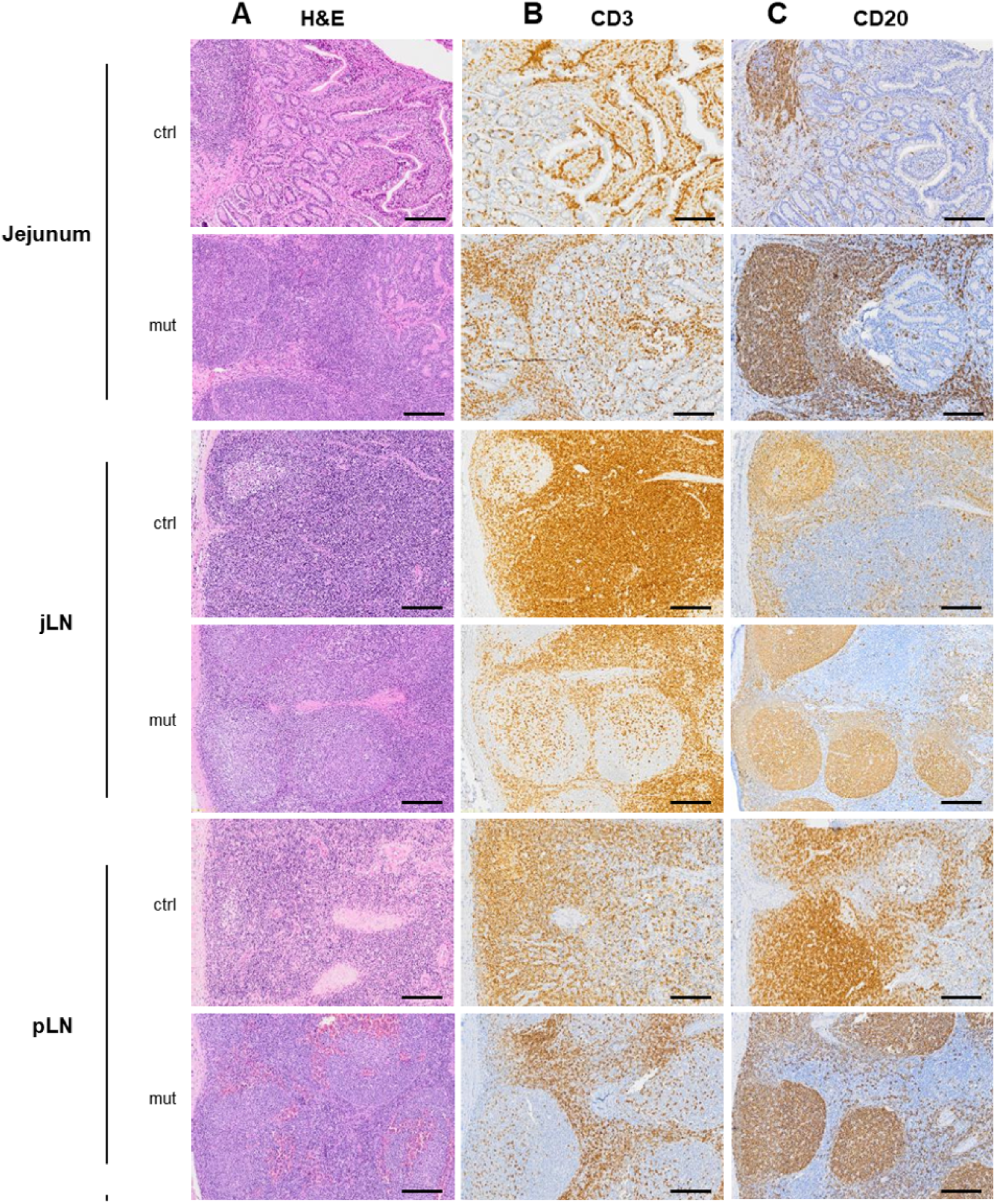
Histopathology of the jejunum, jejunal lymph node (jLN) and peripher<u>al</u> lymph nodes (pLN) of a mutant and a wild-type homozygous animal. (**A**) Haematoxylin and eosin staining of jejunum, jejunal lymph node and peripheral lymph node tissues of a mutant (mut) and a wild type (ctrl). (**B**) CD3 (T lymphocyte marker) and (**C**) CD20 (B lymphocyte marker) immunostaining of the same tissues of the mutant and wild type (ctrl VS mut).

### ITGB7 p.G375S substitution affects the faecal microbiota in homozygous mutant carriers

The systemic and local (intestinal) immune system is exposed to and interacts closely with the microbiota in the digestive tract as the main trigger of its development ^45^. In link with the evident changes in the adaptive compartment of the digestive tract, we investigated the difference in the faecal microbiota composition. The microbiota composition of the faeces was examined in 24 couples of mutants and controls in different commercial farms and represented on a phylum scale (**Figure 5A**). We identified notable differences in the taxonomic composition between the two groups, with some bacterial populations that are in greater proportions in the cases. Indeed, as **Figure 5B** illustrates, there are substantial disparities in alpha-diversity between the two groups (p-value <0.05), as evidenced by all indicators including observed richness, Chao1, and Shannon’s, except for the inverse Simpson index, indicating a disruption in microbial biodiversity in the mutant group. **Figure 5C** illustrates beta-diversity through Bray-Curti’s dissimilarity distances, showing no discernible separation in the microbiota composition between the mutant and control animals. We next interrogated the difference at the Genus level and found significant differences in microbial abundance for *Escherichia/Shigella* (2.00 log-fold change), *Faecalitalea* (1.45 log-fold change), *[Ruminococcus] torques group* (1.06 log-fold change) and unknown genus from *p-2534-18B5 gut group* (–1.63 log-fold change) illustrated in **Figure 5D**. Our findings show a substantial alteration in the microbiota composition in the mutant compared to their wild-type counterparts.

**Figure 5:**
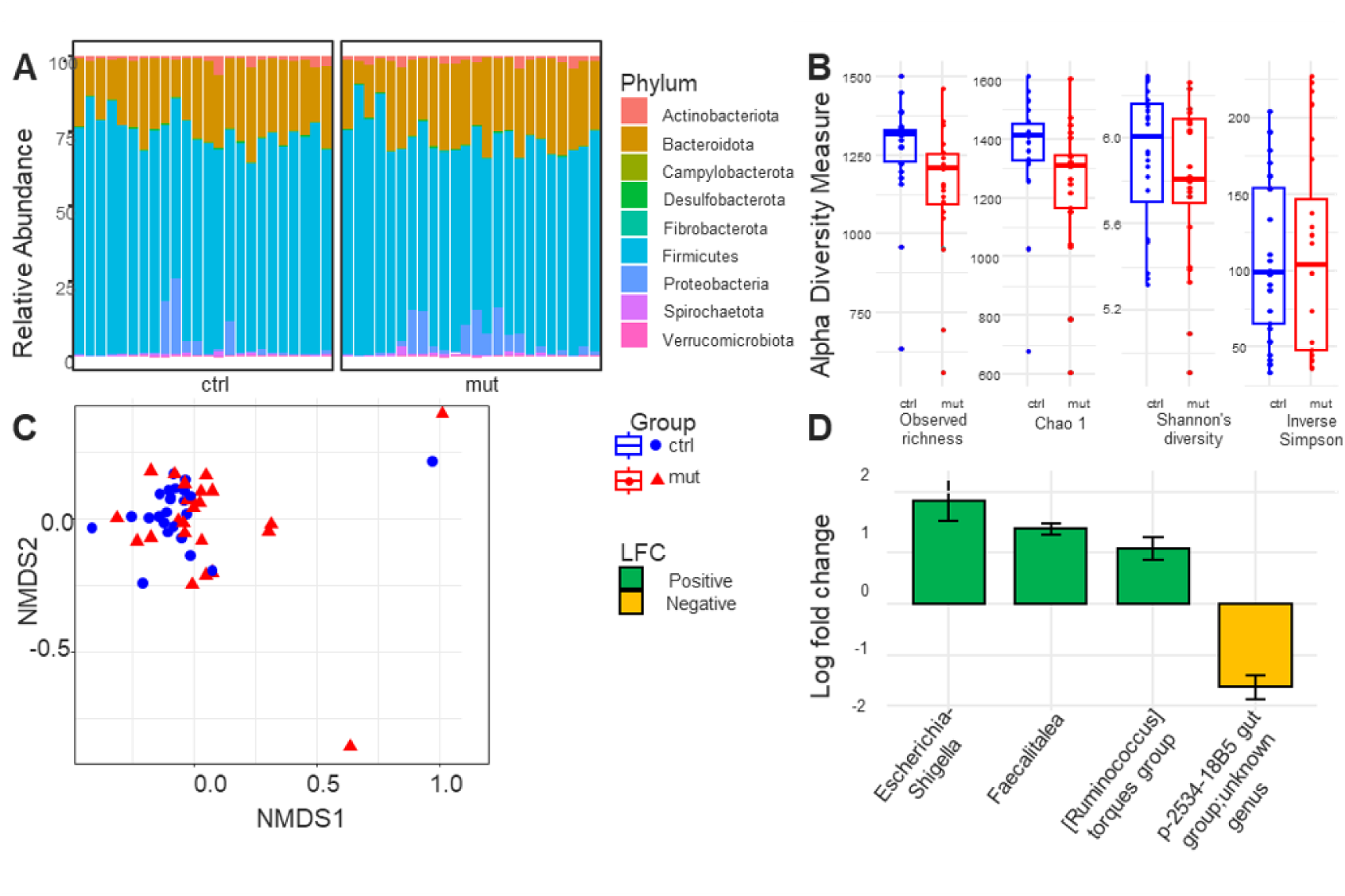
Composition of the faecal microbiome of 24 couples of mutant and control animals in different herds. (**A**) Relative abundance of phyla between controls (ctrl) and mutant (mut) animals. (**B**) Alpha diversity measures, including observed richness, Chao1, Shannon’s diversity index, and inverse Simpson index, comparing controls (ctrl) and mutant (mut) groups. (**C**) Beta diversity, represented by Bray-Curtis dissimilarity distances between controls (blue rounds) and mutant (red triangles) animals. (**D**) Log fold change at the Genus level comparing mutant (mut) to control (ctrl) animals, showing only significant differences.

### Default stature and few blood parameters typify carrier from non-carrier status

The subsequent inquiry sought to ascertain the ease with which homozygote carriers could be identified within the cattle population. Utilising multifactorial analysis (MFA) and principal component analysis (PCA), we examined the influence of genotype and the values of associated uncorrelated biological variables on classification. PCA biplots demonstrated that individuals with identical genotypes formed distinct clusters, thereby substantiating the robustness of genotype-related patterns. Hierarchical clustering further demonstrated that individuals were grouped according to Farm, indicating a potential effect of the environment on the classification process. The sPLS-DA assessment revealed that Principal Component 1 (PC1), accounting for 24.5% of the total variance, is predominantly comprised of lymphocytes, with monocytes, eosinophils, basophils, and globulins exhibiting a negative correlation with the mutant status (increased). Principal Component 2 (PC2) is predominantly composed of farm and haemoglobin, which is inversely correlated with the control group (**Figure 6A**). The sPLS-DA components presented in **Figure 6B** showed a distinct separation of the mutant and control samples. To conclude, the sPLS-DA results demonstrated that immune-related markers (e.g., lymphocytes, eosinophils monocytes, and basophils) played pivotal roles in both components, with farm emerging as a significant secondly positive contributor. These findings underscore the intricate relationships between immune markers and genotype, reflecting the multifaceted biological processes and environmental conditions that underlie the phenotype data. This integrative analysis demonstrates that only a limited set of variables, including lymphocyte count and growth retardation, are effective in differentiating mutants from their controls.

**Figure 6.**
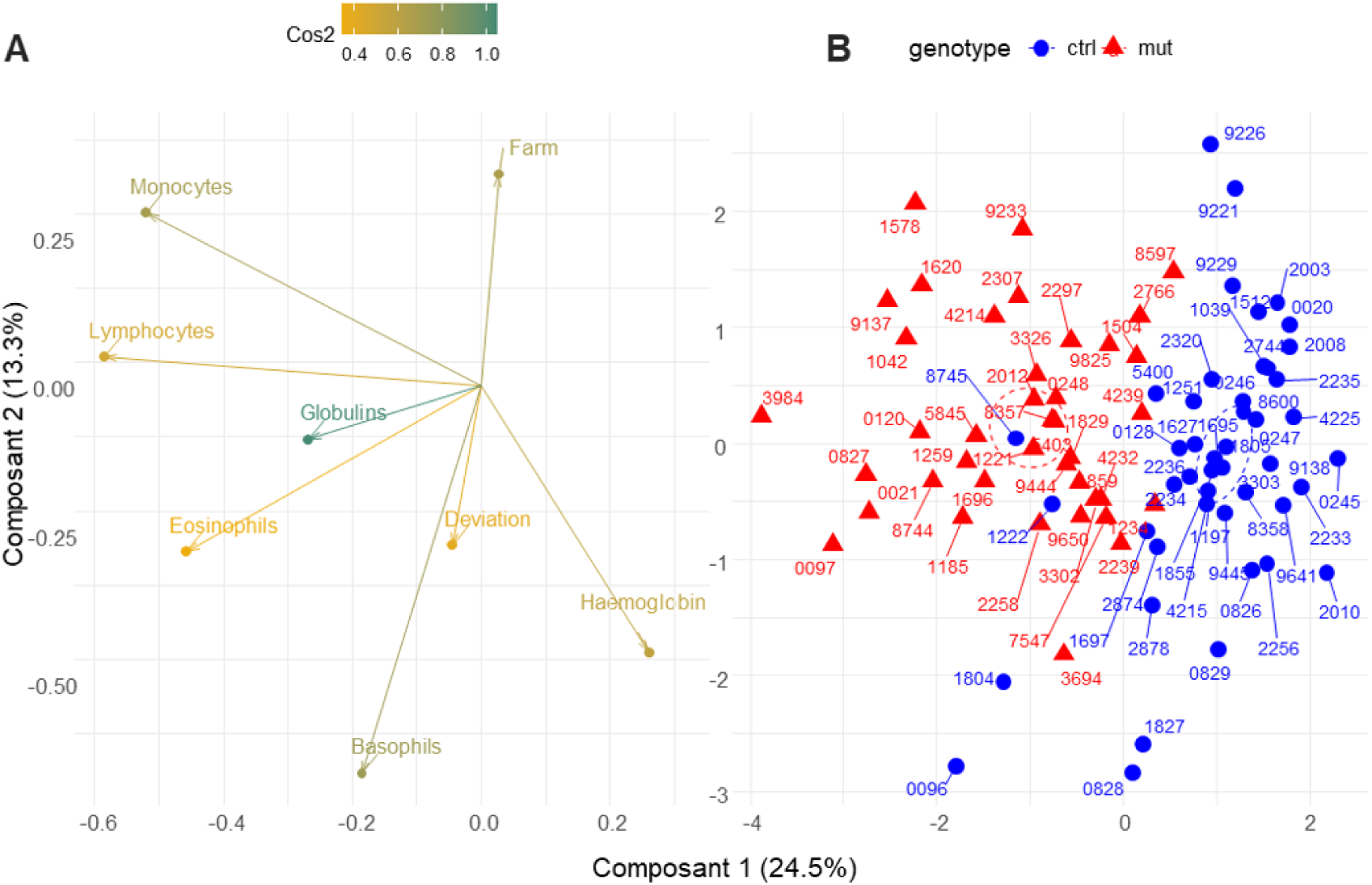
Sparse Partial Least Squares-Discriminant Analysis (PLS-DA) on clinical and biological parameters for identification of mutant animals. (**A**) Representation of variable contributions, with colours indicating their relative importance (cos2). (**B**) sPLS-DA plot for individuals with labels and ellipses with shapes with colour coding based on genetic status with mutant (red triangle), and wild-type (blue round) animals.

## Discussion

Our study aim was to gain insight into the clinical manifestations and underlying mechanisms of BLIRD, a recessive genetic disorder (ITGB7 G375S) recently discovered in Holstein cattle ^19^. We provide essential information for the future management of this genetic disorder in the affected dairy cattle population and interesting observations on the immune system defects associated with a dysfunctional variant of integrin beta7 (ITGB7).

Growth retardation is common in cattle and identification of the underlying cause is a major challenge for cattle practitioners, leading to suboptimal performance and reduced longevity in breeding programs ^46^. In general, poor conditions can be due to malnutrition, parasitism, chronic infections, and less commonly, genetic disorders. Furthermore, environmental factors, including nutrition, housing, infectious disease exposure, and management practices, may further influence the carrier outcome but were out of the scope. Indeed, in our study, animals with the ITGB7 substitution show reduced longevity compared to their controls, bred under the same conditions. Several factors may explain this result. Some animals died prematurely, exhibiting progressive emaciation, poor condition and sudden death at a time of expected exponential growth. First, numerous diseases could be implicated, but according to breeders, no specific illness was detected, and none of the attempted treatments cleared the problem. Second, too low development and insufficient stature at breeding time lead to early culling. The optimal age for AI in dairy heifers is approximately 15 months, corresponding to around 55% of their mature weight ^44^ to ensure they attain sexual maturity and achieve optimal fertility ^47^. However, delaying AI beyond this recommended timeframe results in diminished reproductive efficiency, characterized by prolonged inter-calving intervals, reduced lifetime productivity, and the risk of health complications and exacerbated metabolic disorders. Thus, delayed reproduction leads to extended periods of non-productive life and reduces the sustainability of dairy production ^48^. Third, lower lactation performances for primiparous cows are another reason for reduced productive life by early culling ^49^ ^4^. Conversely, milk quality assessed by milk somatic cell concentrations was not significantly different, suggesting that a predisposition to common infections, like mammary infection and mastitis, cannot be suspected. Moreover, it should be noted that the statistical power of the analyses is limited by the relatively low number of homozygotes available and that the magnitude of the effect may be underestimated because morphological traits are measured during the first lactation and production traits are censored for animals with a lactation length lower. We finally provide clinical descriptions and quantitative parameters that may help detect suspected cases without genetic testing, which is however the ultimate way of confirmation.

In our study, animals with the ITGB7 substitution are smaller and in poorer condition than matched controls. To reduce the impact of confounding variables, the age difference in each couple of control and case in the same herd was maintained at a minimum, typically less than a month in most cases. However, due to their shorter stature, homozygous carriers of the point mutation were frequently raised with younger stock. This resulted in heifers of similar age being located in different areas or eventually absent in small-size herds, thus unavailable for pairing. This discrepancy in age between carriers and controls can be attributed to the choice of controls being younger than mutants. The statistical approach adopted effectively reduced the confounding effect of age, thereby providing a clearer understanding of the genotype effect on several measures, including chest circumference.

Thus, the field analyses have shown how the ITGB7 G375S mutation affects the longevity of Holstein dairy cows, with growth retardation and inappropriateness to breeding in homozygous animals under most herd conditions. No impaired, nor favourable traits were observed in the heterozygotes compared to matched controls suggesting no advantage in heterozygosity. Considering the serious consequences of the mutant variant on the health and welfare of dairy cows, it is recommended to avoid further breeding of carriers, so that the mutation can progressively be eliminated from the cattle population. Digital breeding management can help reduce the incidence of this genetic disorder by allowing early detection and enabling breeders to make informed decisions about mating strategies ^19^.

To get further, the role of the ITGB7 p.G375S default was investigated. The ITGB7 protein is a subunit of the α4β7 LPAM-1 integrin dimer and plays a critical role in the tissue distribution of several immune cell types under homeostatic conditions. Due to the LPAM-1/MAdCAM-1 interaction on intestinal High Endothelial Venules (HEVs) for cell relocation, the mutation could have impaired the animal’s ability to respond to a digestive infection by any pathogen such as viruses, bacteria or helminths. Our observations do not support this hypothesis as no digestive signs such as diarrhoea were reported, and we did not find any gross or microscopic lesions in the intestines of homozygous carriers of the variant that would indicate such a predisposition. However, a few homozygote mutants were necropsied, and an observational bias in the case recruitment cannot be excluded which could have minored the severity of lesions in a less affected category of necropsied animals. Amongst the α4β7 LPAM-1-expressing cells, the homing of CD4 T lymphocytes to the gut-associated lymphoid tissues (GALT) is best described ^21^. In homozygous cattle, the point mutation may disrupt the dimer expression, thereby impairing lymphocyte diapedesis, with a consequent and significant reduction in the number of memory CD4 T cells in the lamina propria, where these cells are typically readily identifiable. This finding suggests that these animals may experience an impaired immune memory response in the digestive tract, and more systemically.

Several interesting and novel observations were made. We report a significant increase in circulating leukocytes, with blood lymphocytes, eosinophils, and monocytes showing significantly higher concentrations in the homozygous mutants compared to the controls.

Several studies have described the expression of ITGB7 at the cell membrane of these cells, indicating that this adhesion molecule may be necessary for their relocation and full activity. In ITGB7 knockout mice ^50^, or following treatment with Vedolizumab, a monoclonal antibody that blocks the interaction between LPAM-1 and MadCAM-1 ^51^; lymphocyte migration to the GALT is impaired, potentially disrupting intestinal homeostasis and immune tolerance. Consistent with these findings, our examination reveals a deficit in LPAM-1 expression in the mutants and precisely on activated/memory CD4+ T cells. However, unlike these studies, we observed an accumulation of those lymphocytes in peripheral blood, which may be explained by the inability of these cells to migrate toward endothelial cells. Furthermore, the lack of migration of various lymphocyte subsets could hinder proper immune surveillance, increasing susceptibility to parasitic infections, as demonstrated in mice infected with *Trichinella spiralis* ^52^. Additionally, in cases of BLIRD, a pronounced lymphocytosis was observed, alongside eosinophilia and monocytosis. Lymphocytosis is a rare condition in cattle, often diagnosed during chronic viral or purulent infections kidney or liver inflammation, as well as bovine leukaemia virus (BLV) characterized by increased B-lymphocytes ^53^, but all herds and animals were free of these infections. Interestingly, the ITGB7 G375S point mutation appears to contribute to lymphocytosis, in the absence of overt viral infections, suggesting that the genetic disorder itself may be associated with a chronic state of immune activation, with accumulation of memory CD4 T lymphocytes in the bloodstream and follicular hyperplasia in the lymph nodes. This is consistent with findings from other studies, which have shown that mutations affecting immune system function, such as those in integrins, can lead to persistent lymphocyte proliferation ^54,55^. The chronic nature of lymphocytosis in these animals, combined with the concurrent eosinophilia, and monocytosis may exacerbate immune dysfunction, highlighting the importance of careful monitoring and management of this genetic anomaly in the cattle population.

Eosinophilia develops due to allergy, inflammatory responses and neoplasm ^56,57^. It’s also associated with parasitic infections ^58^ such as the response to helminths, particularly by gastrointestinal nematodes (e.g., *Ostertagia spp.*, *Cooperia spp.*) and lungworms (e.g., *Dictyocaulus viviparus)* no preferential excretion in the mutants was shown in our study. This may be related to low/no exposure due to no grazing in most farms. Parasitism is not a sound explanation for ill-thrift and growth retardation. Hypersensitivity reactions or fungal infections can trigger a similar immune response, particularly in chronic or granulomatous inflammation, in which eosinophils contribute to tissue repair ^59^. Additionally, autoimmune diseases and certain neoplasms can cause persistent eosinophilia due to abnormal immune system activation and cellular proliferation ^60,61^ but no evidence was collected to support this possibility.

Most mutants presented a low haemoglobin concentration, a common and multifactorial condition, arising from various causes, including poor condition. Several deficiencies for iron or copper ^62^, and chronic inflammatory conditions can suppress marrow red cell production. Parasitic infections by hematophagous pathogens are also a common cause of anaemia ^63^, but again none was present.

An increase in plasma globulins was also noted, which is commonly seen as a result of chronic immune activation, often observed in cattle with persistent infections or ageing. Here, it could be associated with the follicular hyperplasia observed in all the examined lymph nodes, although the exact underlying mechanism has yet to be determined. Lympho-nodal follicular hyperplasia is a reactive benign proliferation affecting the lymphoid cell compartment. It is associated with adaptative B-cell immune tissue in response to many factors including infections, immune-mediated disorders, and certain malignancies. The continuous antigenic challenge comes from the feed and gut microbiota that play a pivotal role in shaping the immune system, particularly in cattle, where the rumen and intestines are home to large and diverse microbial communities ^64^. Diet, environmental conditions, and genetic factors all contribute to the composition of the gut microbiota, which in turn affects the host’s immune system ^65^.

Cendron et al. (2020) examined the faecal microbiomes of Holstein-Friesian and Simmental heifers and lactating cows, identifying significant differences in microbial composition between the two groups within the same herd. The predominant bacterial phyla were *Firmicutes*, *Bacteroidetes*, *Actinobacteria*, and *Proteobacteria*, with variations in bacterial families and genera observed between heifers and cows, suggesting that the faecal microbiome can be influenced by management conditions. While age-related variation could introduce bias in our study, the mutated animals, although slightly older, were on the same diet and at the same physiological stage as their controls, minimizing this factor. In case of immune dysfunction, disruptions in the gut microbiota may exacerbate immune system imbalance ^67^, further compromising health and growth ^68^, such as those with the ITGB7 mutation, Indeed, examination of the gut microbiota’s influence on immune system function is particularly relevant in the context of the ITGB7 mutation, as impaired immune cell migration to the GALT could lead to an altered microbial environment of the digestive tract and reciprocally. This disruption may affect the balance between beneficial and harmful microbiota, further impairing digestion, nutrient absorption, and immune surveillance. In contrast to what has been described in knockout mice where a few Operational Taxonomic Units (OTU) were a little more abundant ^70^, we found a clear difference in the microbiota composition between mutants and their within-farm matched controls. First, α and β diversities were affected, and the abundances of several bacteria phyla and genera were significantly different between the two variant categories. Interestingly, several of these bacteria were previously associated with pathological conditions in Bovidae. The *Ruminococcaceae NK4A214 group* was more abundant in the mutant group as similarly described by Ma et al. (2020), studying the gastrointestinal tract bacterial communities in normal and growth-retarded yaks. Wu et al., (2022) examined the gut microbiota of yaks afflicted by diarrhoea, unveiling that the microbial composition exhibited substantial alterations in the diarrhoea group compared to healthy controls. As in the aforementioned study, we observed a decrease in alpha diversity in the mutant group, alongside an increase in the genus *Escherichia-Shigella*, when comparing the mutant and the control groups. Similarly, in their study, Zhen et al. (2022) found a significant reduction in microbial diversity in diarrheic animals, along with alterations in the composition of the gut microbiota in diarrheic Père David’s deer. Additionally, alteration in the gut microbiota could lead to nutritional deficiencies and further compromise the animal’s feeding behaviour, growth and productivity ^69,73^.

One other possibility that should be considered is the metabolic origin of the poor growth with inadequate assimilation of nutrients eventually complicated by the inability of affected animals to effectively utilize nutrients. Abnormal absorption of nutrients in the intestine is plausible but the lack of structural changes and intestinal chronic inflammation lower the probability. Immune dysfunction often leads to insulin resistance impairing nutrient uptake and utilisation, further contributing to observed weight loss and diminished body condition ^74^. Moreover, some integrin beta7^pos^ T cells regulate the digestive epithelium for nutrient absorption by mitigating GLP-1 availability, rendering *Itgb7* knockout mice metabolically hyperactive ^75^. These mechanisms have not yet been described or even studied in the bovine species.

This study of the BLIRD genetic disorder provides valuable insights into immune defects in Holstein cattle; several limitations and potential biases must however be acknowledged. The willingness of farmers to participate in the survey was an uncontrollable factor affecting the sampling process. The cattle were recruited from a pool of genotyped cattle in France, which may have introduced a bias in the prevalence and symptoms observed in the general population since the AI bull usage is sometimes not the same in high genetic value stocks compared to other commercial farms. The farms participating in the genotyping program have generally more advanced technical and management practices, which could influence the observed outcomes. Out of 40 recruited mutants, only 26 pairs of matched case controls were formed due to the incapacity to find an appropriate control within the herd at the time of the visit. Furthermore, necropsied animals were chosen amongst the group of homozygotes from the epidemiological study and were representative of most clinical presentations. Despite their small number, the fact that the lesions match each other makes us confident that these observations are typical of BLIRD.

BLIRD represents a novel genetic disorder in Holstein cattle, with significant implications for animal health and breeding practices. The present study proposes a novel approach to the investigation of the function of β7 integrin, employing an animal model that allows for the comparison of results across cattle and other species. Furthermore, α4β7-MAdCAM-1 interaction is also a therapeutic target in inflammatory bowel diseases (IBD) in humans and developing more effective therapeutic agents and broader applications beyond is needed. Comparative and translational immunology between species may help understand β7 integrin function in the development and homeostasis of the immune system.

## Acknowledgements

We would like to express our sincere gratitude to the breeders, veterinarians, and agricultural technicians who contributed to this study by granting access to animals and samples.

## Authors’ contributions statement

GF conceived and coordinated the project. LD and GF designed the experiments. AC and FB analyzed survival curves, phenotypic traits, and performances at population level. LD and FC performed the statistical analyses on phenotypes. LG-P and BG conducted laboratory and cytology analyses, including serologies and qPCR. NG, YA-M, AP, LD, and GF performed necropsy examinations and histological analyses. LA analyzed the microbiota. LD and GF wrote the manuscript, with contributions from AC, FB, BG, NG, YA-M, LD, and LA. All authors reviewed and revised the manuscript critically for intellectual content and approved the final version to be published. All authors agree to be accountable for all aspects of the work.

## References

[1] Wiggans GR, Carrillo JA. Genomic selection in United States dairy cattle. Front Genet. 2022;13:994466. doi:10.3389/fgene.2022.994466

[2] Dallago GM, Wade KM, Cue RI, McClure JT, Lacroix R, Pellerin D, et al. Keeping Dairy Cows for Longer: A Critical Literature Review on Dairy Cow Longevity in High Milk-Producing Countries. Animals. 2021;11(3):808. doi:10.3390/ani11030808

[3] Schuster JC, Barkema HW, De Vries A, Kelton DF, Orsel K. Invited review: Academic and applied approach to evaluating longevity in dairy cows. Journal of Dairy Science. 2020;103(12):11008–11024. doi:10.3168/jds.2020-19043

[4] De Vries A. Symposium review: Why revisit dairy cattle productive lifespan? Journal of Dairy Science. 2020;103(4):3838–3845. doi:10.3168/jds.2019-17361

[5] Guinan FL, Wiggans GR, Norman HD, Dürr JW, Cole JB, Van Tassell CP, et al. Changes in genetic trends in US dairy cattle since the implementation of genomic selection. Journal of Dairy Science. 2023;106(2):1110–1129. doi:10.3168/jds.2022-22205

[6] Mugambe J, Ahmed RH, Thaller G, Schmidtmann C. Impact of inbreeding on production, fertility, and health traits in German Holstein dairy cattle utilizing various inbreeding estimators. Journal of Dairy Science. 2024;107(7):4714–4725. doi:10.3168/jds.2023-23728

[7] Robertson a. A theory of limits in artificial selection. Published online March 14, 1960.

[8] Charlesworth B, Charlesworth D. The genetic basis of inbreeding depression. Genet Res. 1999;74(3):329–340. doi:10.1017/S0016672399004152

[9] Daetwyler HD, Villanueva B, Bijma P, Woolliams JA. Inbreeding in genome-wide selection. J Animal Breeding Genetics. 2007;124(6):369–376. doi:10.1111/j.1439-0388.2007.00693.x

[10] Michot P, Chahory S, Marete A, Grohs C, Dagios D, Donzel E, et al. A reverse genetic approach identifies an ancestral frameshift mutation in RP1 causing recessive progressive retinal degeneration in European cattle breeds. Genetics Selection Evolution. 2016;48(1):56. doi:10.1186/s12711-016-0232-y

[11] Reynolds G, Vegh P, Fletcher J, Poyner EFM, Stephenson E, Goh I, et al. Developmental cell programs are co-opted in inflammatory skin disease. Science. 2021;371(6527):eaba6500. doi:10.1126/science.aba6500

[12] Agerholm JS, McEvoy F, Arnbjerg J. Brachyspina Syndrome in a Holstein Calf. J VET Diagn Invest. 2006;18(4):418–422. doi:10.1177/104063870601800421

[13] Charlier J, Höglund J, Morgan ER, Geldhof P, Vercruysse J, Claerebout E. Biology and Epidemiology of Gastrointestinal Nematodes in Cattle. Veterinary Clinics of North America: Food Animal Practice. 2020;36(1):1–15. doi:10.1016/j.cvfa.2019.11.001

[14] Häfliger IM. Forward vs. reverse genetics: a bovine perspective based on visible and hidden phenotypes of inherited disorders. Published online January 17, 2022. doi:10.48549/3486

[15] Shanks RD, Dombrowski DB, Harpestad GW, Robinson JL. Inheritance of UMP synthase in dairy cattle. Journal of Heredity. 1984;75(5):337–340. doi:10.1093/oxfordjournals.jhered.a109951

[16] Shuster DE, Kehrli ME, Ackermann MR, Gilbert RO. Identification and prevalence of a genetic defect that causes leukocyte adhesion deficiency in Holstein cattle. Proc Natl Acad Sci USA. 1992;89(19):9225–9229. doi:10.1073/pnas.89.19.9225

[17] Fritz S, Capitan A, Djari A, Rodriguez SC, Barbat A, Baur A, et al. Detection of Haplotypes Associated with Prenatal Death in Dairy Cattle and Identification of Deleterious Mutations in GART, SHBG and SLC37A2. Veitia RA, ed. PLoS ONE. 2013;8(6):e65550. doi:10.1371/journal.pone.0065550

[18] VanRaden PM, Olson KM, Null DJ, Hutchison JL. Harmful recessive effects on fertility detected by absence of homozygous haplotypes. Journal of Dairy Science. 2011;94(12):6153–6161. doi:10.3168/jds.2011-4624

[19] Besnard F, Guintard A, Grohs C, Guzylack-Piriou L, Cano M, Escouflaire C, et al. Massive detection of cryptic recessive genetic defects in dairy cattle mining millions of life histories. Genome Biol. 2024;25(1):248. doi:10.1186/s13059-024-03384-7

[20] Rüegg C, Postigo A, Sikorski E, Butcher E, Pytela R, Erle D. Role of integrin alpha 4 beta 7/alpha 4 beta P in lymphocyte adherence to fibronectin and VCAM-1 and in homotypic cell clustering. The Journal of cell biology. 1992;117(1):179–189. doi:10.1083/jcb.117.1.179

[21] Gorfu G, Jesús RN, Ley K. Role of β7 integrins in intestinal lymphocyte homing and retention. Published online September 2009.

[22] Berlin C, Berg EL, Briskin MJ, Andrew DP, Kilshaw PJ, Holzmann B, et al. α4β7 integrin mediates lymphocyte binding to the mucosal vascular addressin MAdCAM-1. Cell. 1993;74(1):185–195. doi:10.1016/0092-8674(93)90305-A

[23] Cimbro R, Vassena L, Arthos J, Cicala C, Kehrl JH, Park C, et al. IL-7 induces expression and activation of integrin α4β7 promoting naive T-cell homing to the intestinal mucosa. Blood. 2012;120(13):2610–2619. doi:10.1182/blood-2012-06-434779

[24] Andrew DP, Rott LS, Kilshaw PJ, Butcher EC. Distribution of α4β7 and αEβ7 integrins on thymocytes, intestinal epithelial lymphocytes and peripheral lymphocytes. European Journal of Immunology. 1996;26(4):897–905. doi:10.1002/eji.1830260427

[25] Keir ME, Fuh F, Ichikawa R, Acres M, Hackney JA, Hulme G, et al. Regulation and Role of αE Integrin and Gut Homing Integrins in Migration and Retention of Intestinal Lymphocytes during Inflammatory Bowel Disease. The Journal of Immunology. 2021;207(9):2245–2254. doi:10.4049/jimmunol.2100220

[26] Schleier L, Wiendl M, Heidbreder K, Binder MT, Atreya R, Rath T, et al. Non-classical monocyte homing to the gut via α4β7 integrin mediates macrophage-dependent intestinal wound healing. Gut. 2020;69(2):252–263. doi:10.1136/gutjnl-2018-316772

[27] Walsh. Integrin a a4b b7 mediates human eosinophil interaction with MAdCAM-1, VCAM-1 and fibronectin. Published online 1996.

[28] Shouval DS. α4β7 expression guides B cells to front lines of defense in the gut. Mucosal Immunology. 2022;15(2):192–194. doi:10.1038/s41385-021-00476-6

[29] Wagner SD. Somatic Hypermutation Of Immunoglobulin Genes. Published online 1996.

[30] Feagan BG, Rutgeerts P, Sands BE, Hanauer S, Colombel JF, Sandborn WJ, et al. Vedolizumab as Induction and Maintenance Therapy for Ulcerative Colitis. N Engl J Med. 2013;369(8):699–710. doi:10.1056/NEJMoa1215734

[31] Misztal I, Lourenco D, Aguilar I, Legarra A, Vitezica Z. Manual for BLUPF90 family of programs. Published online May 2014.

[32] Benjamini Y, Hochberg Y. Controlling the False Discovery Rate: A Practical and Powerful Approach to Multiple Testing. Journal of the Royal Statistical Society Series B: Statistical Methodology. 1995;57(1):289–300. doi:10.1111/j.2517-6161.1995.tb02031.x

[33] Jourdain J, Barasc H, Faraut T, Calgaro A, Bonnet N, Marcuzzo C, et al. Large-scale detection and characterization of interchromosomal rearrangements in normozoospermic bulls using massive genotype and phenotype data sets. Genome Res. 2023;33(6):957–971. doi:10.1101/gr.277787.123

[34] Grebert M, Granat F, Braun J, Leroy Q, Bourgès-Abella N, Trumel C. Validation of the Sysmex XN-V hematology analyzer for canine specimens. Veterinary Clinical Pathol. 2021;50(2):184–197. doi:10.1111/vcp.12936

[35] Baltrušis P, Halvarsson P, Höglund J. Molecular detection of two major gastrointestinal parasite genera in cattle using a novel droplet digital PCR approach. Parasitol Res. 2019;118(10):2901–2907. doi:10.1007/s00436-019-06414-7

[36] R Core Team. R: A language and environment for statistical computing. Published online 2023. https://www.r-project.org

[37] Brooks, Mollie E. and Kristensen, Kasper and Benthem, Koen J. van and Magnusson, Arni and Berg, Casper W. and Nielsen, Anders and Skaug, Hans J. and Mächler, Martin and Bolker, Benjamin M. glmmTMB Balances Speed and Flexibility Among Packages for Zero-inflated Generalized Linear Mixed Modeling. Published online 2017:378–400.

[38] V. Lenth R. emmeans: Estimated Marginal Means, aka Least-Squares Means. Published online 2024. https://CRAN.R-project.org/package=emmeans

[39] Lê S, Josse J, Husson F. FactoMineR: An R Package for Multivariate Analysis. J Stat Soft. 2008;25(1). doi:10.18637/jss.v025.i01

[40] Rohart F, Gautier B, Singh A, Lê Cao KA. mixOmics: An R package for ‘omics feature selection and multiple data integration. Schneidman D, ed. PLoS Comput Biol. 2017;13(11):e1005752. doi:10.1371/journal.pcbi.1005752

[41] Oksanen, J., Blanchet, F. G., Friendly, M., Kindt, R., Legendre, P., McGlinn, D., Minchin, P. R., O’Hara, R. B., Simpson, G. L., Solymos, P., Stevens, M. H. H., Szoecs, E., & Wagner, H. vegan: Community Ecology Package. Published online 2022.

[42] Lin, D. Y., Eggesbø, M., & Peddada, S. D. An improved bias correction method for the analysis of microbiome composition data. Published online 2022. 10.3389/fmicb.2022.815366

[43] Lin, D. Y., & Peddada, S. D. Analysis of compositions of microbiomes with bias correction (ANCOM-BC). Published online 2020. 10.1038/s41467-020-15560-z.

[44] Smith J. Heifer growth and economics: target growth. Published online 2007.

[45] Molloy MJ, Bouladoux N, Belkaid Y. Intestinal microbiota: Shaping local and systemic immune responses. Seminars in Immunology. 2012;24(1):58–66. doi:10.1016/j.smim.2011.11.008

[46] Heinrichs AJ, Zanton GI, Lascano GJ, Jones CM. A 100-Year Review: A century of dairy heifer research. Journal of Dairy Science. 2017;100(12):10173–10188. doi:10.3168/jds.2017-12998

[47] Le Cozler Y, Lollivier V, Lacasse P, Disenhaus C. Rearing strategy and optimizing first-calving targets in dairy heifers: a review. Animal. 2008;2(9):1393–1404. doi:10.1017/S1751731108002498

[48] Wathes DC, Pollott GE, Johnson KF, Richardson H, Cooke JS. Heifer fertility and carry over consequences for life time production in dairy and beef cattle. Animal. 2014;8:91–104. doi:10.1017/S1751731114000755

[49] De Vries A, Marcondes MI. Review: Overview of factors affecting productive lifespan of dairy cows. Animal. 2020;14:s155–s164. doi:10.1017/S1751731119003264

[50] Wagner N, Löhler J, Kunkel EJ, Ley K, Leung E, Krissansen G, et al. Critical role for β7 integrins in formation of the gut-associated lymphoid tissue. Nature. 1996;382(6589):366–370. doi:10.1038/382366a0

[51] Canales-Herrerias P, Uzzan M, Seki A, Czepielewski RS, Verstockt B, Livanos AE, et al. Gut-associated lymphoid tissue attrition associates with response to anti-α4β7 therapy in ulcerative colitis. Sci Immunol. 2024;9(94):eadg7549. doi:10.1126/sciimmunol.adg7549

[52] Song Y, Xu J, Wang X, Yang Y, Bai X, Pang J, et al. Regulation of host immune cells and cytokine production induced by *Trichinella spiralis* infection. Parasite. 2019;26:74. doi:10.1051/parasite/2019074

[53] Abramowicz B, Kurek Ł, Lutnicki K. Haematology in the early diagnosis of cattle diseases – a review. Vet arhiv. 2019;89(4):579–590. doi:10.24099/vet.arhiv.0700

[54] Calderwood DA, Campbell ID, Critchley DR. Talins and kindlins: partners in integrin-mediated adhesion. Nat Rev Mol Cell Biol. 2013;14(8):503–517. doi:10.1038/nrm3624

[55] Moser M, Legate KR, Zent R, Fässler R. The Tail of Integrins, Talin, and Kindlins. Science. 2009;324(5929):895–899. doi:10.1126/science.1163865

[56] Morales-Alvarez MC. Nephrotoxicity of Antimicrobials and Antibiotics. Advances in Chronic Kidney Disease. 2020;27(1):31–37. doi:10.1053/j.ackd.2019.08.001

[57] Ramirez GA, Yacoub MR, Ripa M, Mannina D, Cariddi A, Saporiti N, et al. Eosinophils from Physiology to Disease: A Comprehensive Review. BioMed Research International. 2018;2018:1–28. doi:10.1155/2018/9095275

[58] Claerebout E, Vercruysse J. The immune response and the evaluation of acquired immunity against gastrointestinal nematodes in cattle: a review. Parasitology. 2000;120(7):25–42. doi:10.1017/S0031182099005776

[59] Coden ME, Berdnikovs S. Eosinophils in wound healing and epithelial remodeling: Is coagulation a missing link? Journal of Leukocyte Biology. 2020;108(1):93–103. doi:10.1002/JLB.3MR0120-390R

[60] Diny NL, Rose NR, Čiháková D. Eosinophils in Autoimmune Diseases. Front Immunol. 2017;8:484. doi:10.3389/fimmu.2017.00484

[61] Shomali W, Gotlib J. World Health Organization and International Consensus Classification of eosinophilic disorders: 2024 update on diagnosis, risk stratification, and management. American J Hematol. 2024;99(5):946–968. doi:10.1002/ajh.27287

[62] Underwood EJ, Suttle NF. The Mineral Nutrition of Livestock. 3rd ed. CABI Publishing; 1999. doi:10.1079/9780851991283.0000

[63] Constable PD, Blood DC, Radostits OM. Veterinary Medicine: A Textbook of the Diseases of Cattle, Horses, Sheep, Pigs, and Goats. 11th edition. Elsevier; 2017.

[64] Keum GB, Pandey S, Kim ES, Doo H, Kwak J, Ryu S, et al. Understanding the Diversity and Roles of the Ruminal Microbiome. J Microbiol. 2024;62(3):217–230. doi:10.1007/s12275-024-00121-4

[65] Jami E, Mizrahi I. Composition and Similarity of Bovine Rumen Microbiota across Individual Animals. López-García P, ed. PLoS ONE. 2012;7(3):e33306. doi:10.1371/journal.pone.0033306

[66] Cendron F, Niero G, Carlino G, Penasa M, Cassandro M. Characterizing the fecal bacteria and archaea community of heifers and lactating cows through 16S rRNA next-generation sequencing. J Appl Genetics. 2020;61(4):593–605. doi:10.1007/s13353-020-00575-3

[67] Jiao Y, Wu L, Huntington ND, Zhang X. Crosstalk Between Gut Microbiota and Innate Immunity and Its Implication in Autoimmune Diseases. Front Immunol. 2020;11:282. doi:10.3389/fimmu.2020.00282

[68] Xu Q, Qiao Q, Gao Y, Hou J, Hu M, Du Y, et al. Gut Microbiota and Their Role in Health and Metabolic Disease of Dairy Cow. Front Nutr. 2021;8:701511. doi:10.3389/fnut.2021.701511

[69] Clemmons BA, Voy BH, Myer PR. Altering the Gut Microbiome of Cattle: Considerations of Host-Microbiome Interactions for Persistent Microbiome Manipulation. Microb Ecol. 2019;77(2):523–536. doi:10.1007/s00248-018-1234-9

[70] Babbar A, Hitch TCA, Pabst O, Clavel T, Hübel J, Eswaran S, et al. The Compromised Mucosal Immune System of β7 Integrin-Deficient Mice Has Only Minor Effects on the Fecal Microbiota in Homeostasis. Front Microbiol. 2019;10:2284. doi:10.3389/fmicb.2019.02284

[71] Ma J, Zhu Y, Wang Z, Yu X, Hu R, Wang X, et al. Comparing the Bacterial Community in the Gastrointestinal Tracts Between Growth-Retarded and Normal Yaks on the Qinghai–Tibetan Plateau. Front Microbiol. 2020;11:600516. doi:10.3389/fmicb.2020.600516

[72] Wu ZL, Wei R, Tan X, Yang D, Liu D, Zhang J, et al. Characterization of gut microbiota dysbiosis of diarrheic adult yaks through 16S rRNA gene sequences. Front Vet Sci. 2022;9:946906. doi:10.3389/fvets.2022.946906

[74] Welch CB, Ryman VE, Pringle TD, Lourenco JM. Utilizing the Gastrointestinal Microbiota to Modulate Cattle Health through the Microbiome-Gut-Organ Axes. Microorganisms. 2022;10(7):1391. doi:10.3390/microorganisms10071391

[76] Ingvartsen KL, Moyes K. Nutrition, immune function and health of dairy cattle. Animal. 2013;7:112–122. doi:10.1017/S175173111200170X

[77] He S, Kahles F, Rattik S, Nairz M, McAlpine CS, Anzai A, et al. Gut intraepithelial T cells calibrate metabolism and accelerate cardiovascular disease. Nature. 2019;566(7742):115–119. doi:10.1038/s41586-018-0849-9

